# Structure-Based Computational Analysis of Interactions between Insulin Receptor and Insulin Inhibitory Receptor

**DOI:** 10.1101/2024.09.06.611694

**Authors:** Victor Li, Yinghao Wu

## Abstract

The recently discovered insulin inhibitory receptor (inceptor) plays a crucial role in insulin resistance and diabetes by reducing the insulin receptor count on cell membranes, resulting in higher blood glucose levels and decreased insulin sensitivity. Therefore, understanding the mechanism of how the inceptor insulin receptor complex interacts is exceedingly important. This study uses computational drug discovery to inhibit this interaction. Initially, we employed AlphaFold-Multimer to model the inceptor-insulin receptor protein complex and subsequently identified specific inceptor residues likely involved in binding to the insulin receptor. Through virtual screening, thousands of potential small molecules were found to bind to the inceptor, and 10 with the highest probability were chosen for docking. Beta-L-fucose, beta-D-fucose, and alpha-L-fucose showed the most promising binding energies, meaning these three small molecules can effectively interrupt the binding between the complex. We also computationally mutated the binding site of the insulin receptor and calculated the change in binding energy of the inceptor insulin receptor complex, the most dramatic being a 0.4 kcal mol^-1 change when Arginine mutated to Tryptophan at residue 926. Our study suggests that the mutations led to disease primarily due to the change in interactions of the inceptor insulin receptor complex.

## Introduction

Recently, in 2021, the insulin-inhibitory receptor (inceptor) was discovered to be a major player in patients with insulin resistance and Diabetes [1]. Diabetes is a chronic condition characterized by high blood glucose levels which can lead to a myriad of health complications like cardiovascular disease, nerve damage, kidney failure, and more [2]. Insulin is a hormone produced by the pancreas that binds to insulin receptors causing a cascade of interactions that eventually lead to GLUT-4 Channels opening within cells. When the GLUT-4 Channels open, glucose in the bloodstream enters the cell effectively lowering the glucose levels in the bloodstream and regulating blood glucose levels [3]. However, the inceptor molecule has been observed to bind to insulin receptors on the cell’s surface directly, causing endocytosis where the cell absorbs the insulin receptor inside the cytoplasm (**Figure 1**). This lowers the number of insulin receptors on the cell’s surface, causing the GLUT-4 channels to open less frequently, eventually leading to less controlled blood glucose levels within the bloodstream. Therefore, it is important to study the interactions between the inceptor and insulin receptor.

**Figure 1:**
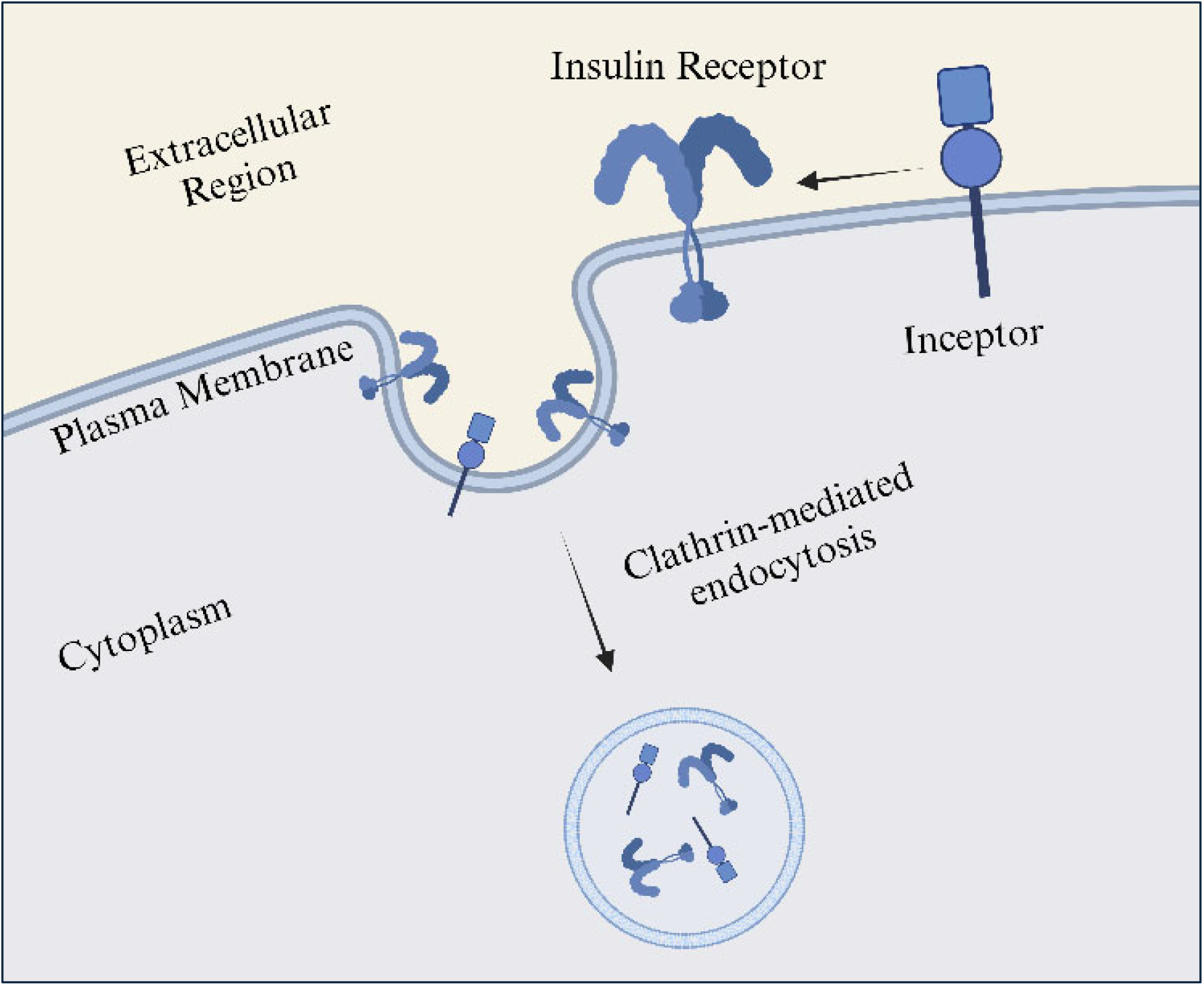
This image shows the interaction between the inceptor and insulin receptors and how they bind to increase insulin resistance

This study applies a computational modeling framework that is widely applicable and important in studying protein-protein interactions and drug discovery. Proteins rarely act alone in a cell but rather function through interactions with other macromolecules, making understanding these interactions on an atomic level fundamental to understanding protein interactions [9]. It is also important to understand proteins on a structural level because proteins with different sequence information can have similar functional structures, that is, different amino acid sequences can show similar folding trends in 3D space and structure is more conserved than sequence. This study accounts for the aforementioned concepts by taking a multi-faceted approach on the cellular and structural levels to inhibit the binding of the inceptor insulin receptor complex. Using AlphaFold Multimer, virtual screening, molecular docking, and mutations, we were able to pinpoint the binding site of the complex along with 10 potential small molecules to prevent binding. We also found 5 mutations that lowered the binding energy of the complex implicating the computational approach.

## Methods

First, we filtered out the regions from the inceptor and insulin receptors that were not related to the interaction. Since these molecules bind on the cell surface, we filtered out the residues in the transmembrane and cytoplasmic regions. We used Uniprot, a web-based database containing protein sequence information, to find the exact residue positions. We filtered out the residue sequence for the inceptor to only include positions 42-910. We filtered the residue sequence for the insulin receptor to only include positions 28-956. Next, we used AlphaFold Multimer to model the structure of the complex formed between the two molecules 5 times for accuracy (**Figure 2**) [4].

**Figure 2:**
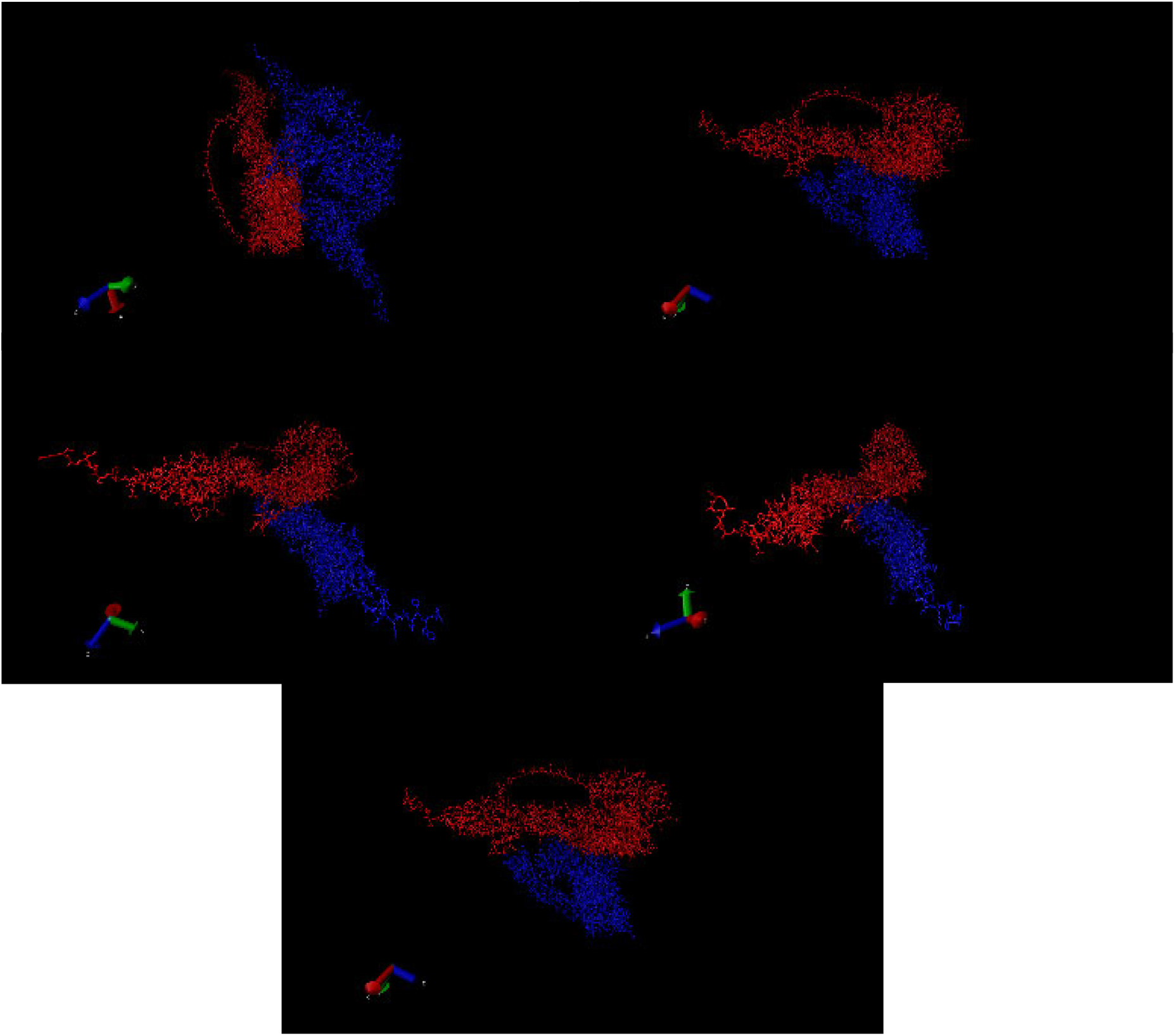
These five models show the interaction between the filtered inceptor and the filtered insulin receptor simulated by AlphaFold multimer. Each model is a separate simulation done for accuracy. The red molecule represents the insulin receptor, and the blue molecule represents the inceptor.

After receiving the modeling results, we converted the simulation file from PDB formatting to an Excel file with information on the cartesian coordinates of each atom, atom number, atom type, residue number, residue type, occupancy, and beta factor of the protein complex. To locate the inceptor’s binding site, we needed to find atoms from the inceptor close to the insulin receptor which would suggest interaction. Since the carbon alpha atom is centrally located within each residue, we needed to determine the distances between these atoms in the inceptor and the insulin receptor within the complex to identify the inceptor’s binding site. So, we first filtered out all non-carbon alpha atoms and wrote a program with Apache POI in Java to find the residues within 10 Angstroms of each other in the complex. We added an “interest” column to indicate if the residue was within 10 Angstroms of another and cross-referenced all 5 models to develop a probability table (**Figure 3**) depicting the residues’ probability of being within the binding site of interest (**Figure 4**). We picked only the residues with a 100% predicted rate across all 5 models to dock small molecules onto. We then used the FindSITE Comb simulation software for virtual screening to generate a list of small molecules that could potentially bind to the inceptor [5]. Finally, we used MTI Autodock, a molecular docking software, to bind 10 of the highest probability small molecules with the specific binding site found earlier [6].

**Figure 3:**
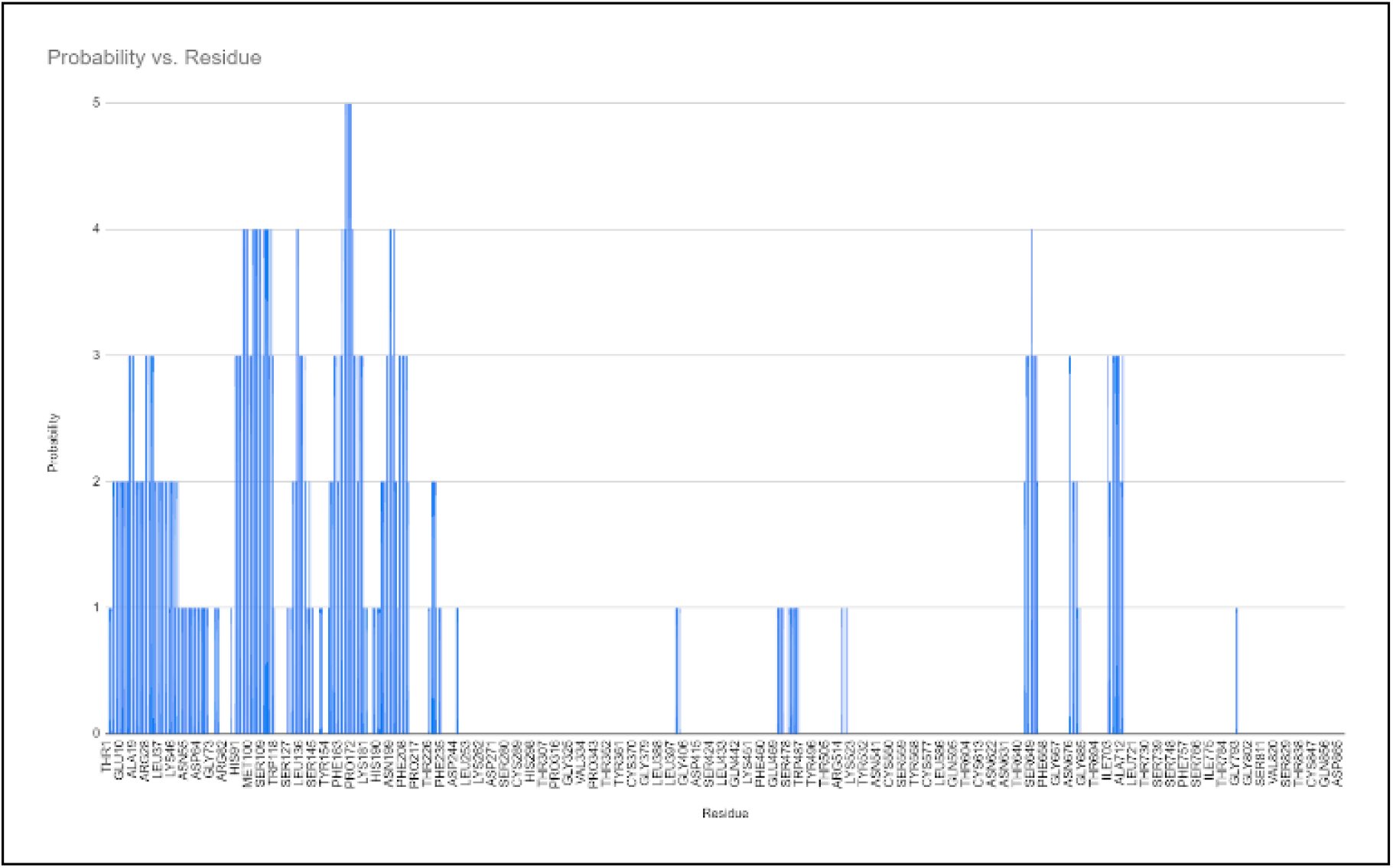
The graph “Probability vs. Residue” shows the likelihood of specific residues within the binding site of the inceptor based on five modeled structures of the inceptor-insulin receptor complex. Each bar represents a residue, with the height indicating the probability of that residue’s involvement in the binding site.

**Figure 4:**
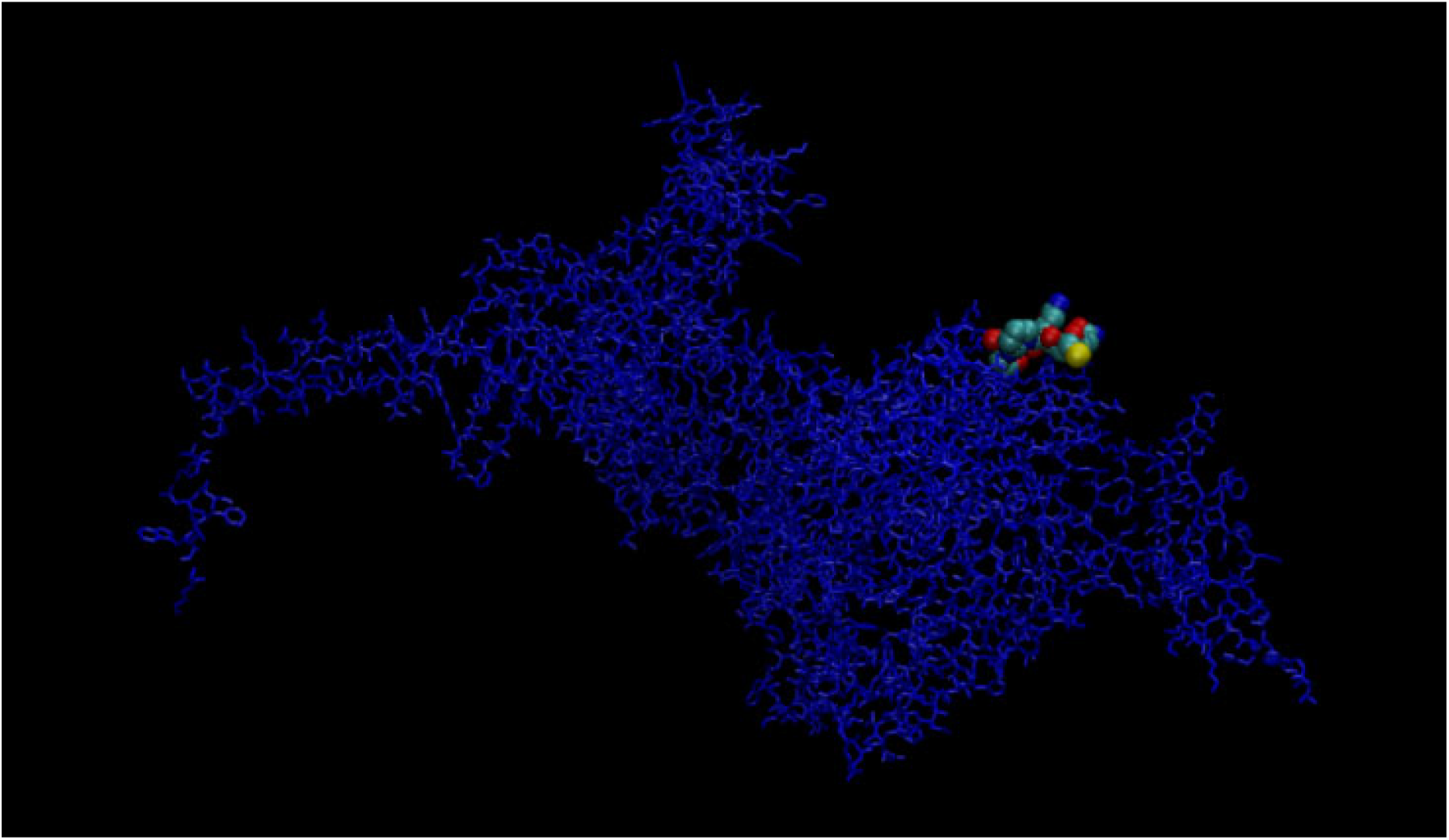
This image shows the inceptor molecule with the binding site that was discovered highlighted as van-der-waal spheres.

After finishing work on the inceptor, we extended the Apache POI program to find the insulin receptor’s binding site using the same method. We also modified the program to keep track of which specific models out of the 5 were predicted to be residue-binding sites. Using Uniprot again, we found the residue numbers with a discovered mutation. We compared the discovered mutation’s residue number with residues that were predicted to be a part of the binding site in any model. When a mutation was found to be within the predicted binding site of the insulin receptor, we modified a duplicate PDB file for the specific model(s) (indicated by the probability program) to contain the mutation without the side chains. To find the effects of the mutation across the entire protein-complex model (including the side chains), we used FASPR, a method for structural modeling of protein side-chain conformations, to accurately predict the whole effect of the mutation [7]. To gauge the effectiveness of the mutations in stopping interactions between the inceptor and insulin receptor we used Prodigy, a web-based server allowing for the prediction of binding energies between biological compounds [8].

## Results and Discussions

We found 10 small molecules most likely to prevent the interaction between the inceptor and insulin receptors (**Table 1**). Out of the 10, the top candidates, beta-L-fucose, beta-D-fucose, and alpha-L-fucose showed the most promising binding energies. These levels of binding energy indicate that these 3 small molecules (**Figure 5**) may have the strongest and most stable bonds with the binding site of the inceptor. The strong binding affinity of these small molecules (particularly beta-L-fucose) implies that they could effectively bind onto the binding site of the inceptor, thereby hindering its interaction with the insulin receptors and preventing insulin resistance (**Figure 6**).

**Table 1:** Ten small molecules with the highest probability of binding onto the binding site of the inceptor in order from highest binding energy to the lowest after docking.

| Molecule Name | Binding Energy ▲ |
| --- | --- |
| beta-L-fucose | -5.97 |
| beta-D-fucose | -5.96 |
| alpha-L-fucose | -5.89 |
| alpha-D-quinovopyranose | -5.84 |
| alpha-D-Fucopyranose | -5.74 |
| alpha-D-quinovopyranose | -5.68 |
| Beta-D-Fructopyranose | -5.44 |
| 4-Deoxy-Alpha-D-Glucose | -5.34 |
| Beta-D-Glucose | -5.04 |
| D-Allopyranose | -4.89 |

**Figure 5:**
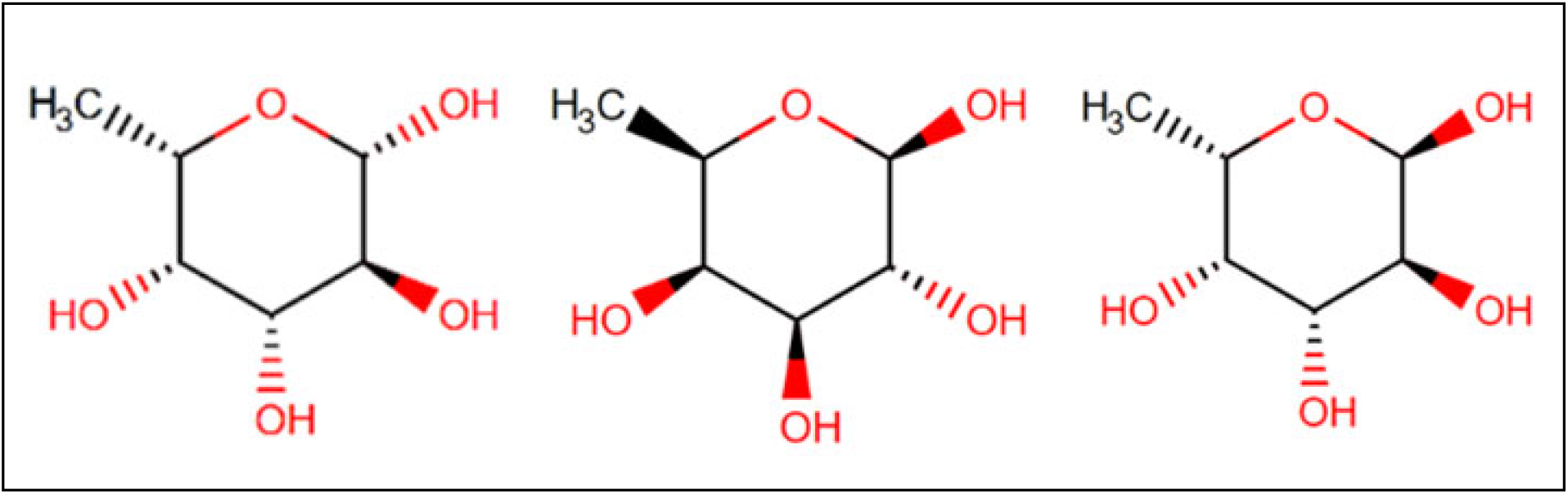
These compounds (Beta-L-Fucose, Beta-D-Fucose, Alpha-L-Fucose from left to right) belong to the class of organic compounds known as hexoses. These are monosaccharides in which the sugar unit is a six-carbon containing moeity. Most of these compounds naturally occur in many biological compounds like the cell surface.

**Figure 6:**
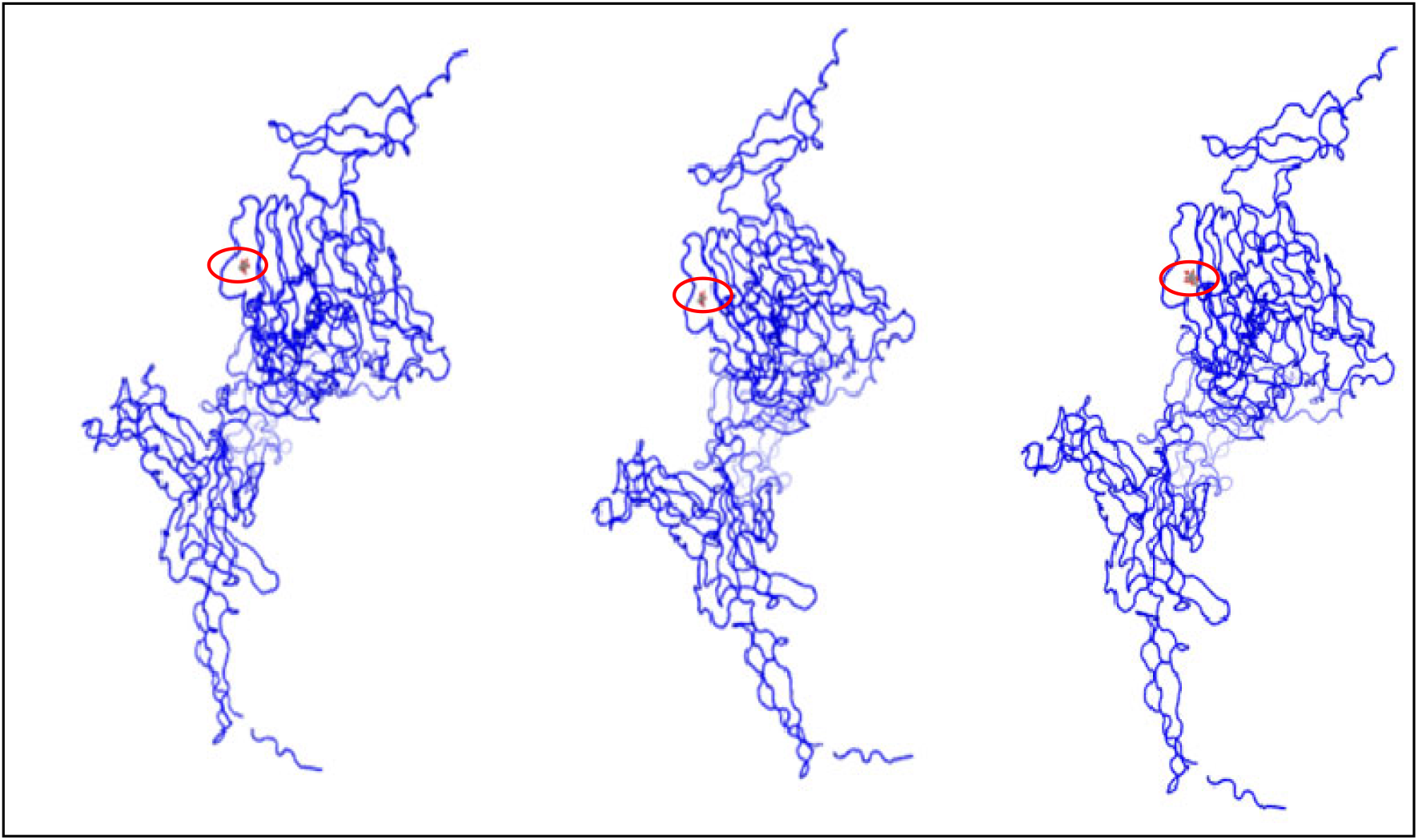
The three highest binding energy small molecules docked onto the inceptor molecule (Beta-L-Fucose, Beta-D-Fucose, Alpha-L-Fucose from left to right).

Using FASPR and Prodigy, we calculated the effect of each mutation on the binding site of the inceptor insulin receptor complex (**Table 2**). We found that the mutations varied in their impact on the mean change in Gibbs free energy (ΔΔG), which reflects the complex’s binding affinity and stability. The data indicates that the Arginine to Tryptophan (R>W) mutation at residue 926 and the Serine to Isoleucine (S>I) mutation at residue 835 have the highest positive and negative impacts on ΔΔG, respectively. This shows that certain mutations can enhance or weaken the binding affinity of the insulin receptor, providing potential targets for modifying the inceptor’s interaction with insulin receptors

**Table 2:** This table shows the type of mutation that occurs and its effects on the binding energy of the complex sorted from greatest weakening impact to greatest strengthening impact. The table also depicts the residue at which the mutations occur and possible diseases they may lead to.

| Residue Number | Mutation | Mean $\Delta\Delta G$ ▼ | Disease Relation |
| --- | --- | --- | --- |
| 926 | R>W | 0.4 | LEPRCH |
| 762 | R>S | 0.2 | IRAN Type A |
| 874 | P>L | 0.1 | RMS |
| 489 | N>D | 0.033 | IRAN Type A |
| 487 | K>E | 0.025 | LEPRCH |
| 458 | N>D | 0 | LEPRCH |
| 878 | N>S | 0 | RMS |
| 937 | T>M | 0 | LEPRCH |
| 842 | A>V | -0.1 | RMS & LEPRCH |
| 818 | Y>C | -0.2 | LEPRCH |
| 350 | S>L | -0.3 | RMS & LEPRCH |
| 925 | I>T | -0.6 | LEPRCH |
| 835 | S>I | -0.75 | RMS & LEPRCH |

This study could only look at the 10 most probable candidates with binding probabilities of 95.0923% of the 6507 discovered due to the limitations of working with public servers. 6497 of the 6507 small molecules identified were unchecked in the docking software due to time and server limitations even if the binding probabilities were around 50% for the majority of the small molecules identified.

The mutations discovered all lead to at least one of three diseases: Insulin-resistant diabetes mellitus with acanthosis nigricans type A (IRAN type A), Leprechaunism (LEPRCH), and Rabson-Mendenhall syndrome (RMS). All three of these conditions are characterized as rare genetic disorders with properties of insulin resistance. IRAN type A typically presents with acanthosis nigricans, a condition marked by dark velvety patches of skin [10]. Leprechaunism, otherwise known as Donohue syndrome, is characterized by severe mental impairment, growth retardation, and an elfin facial appearance, which usually leads to early infant death [11]. Rabson-Mendenhall syndrome has similar symptoms including growth retardation, mental precocity, insulin resistance, and senile-looking faces [12].

Future studies could work to simulate the interactions of more of the 6507 small molecules identified by the virtual screening process. Future studies can also further verify the viability of the small molecules by simulating interactions of the inceptor insulin receptor complex with the small molecule at the binding site. Finally, if more simulations verify these small molecules, we can test them on cells and rats to further gauge their effectiveness as a drug.

## Conclusion

In summary, this study successfully applies a structure-based computational approach to analyze and potentially disrupt the inceptor and insulin inhibitory receptor from binding. The computational framework implemented in this paper reveals the potential for identifying novel therapeutic strategies to combat insulin resistance and diabetes. This unique structure-based approach can be applied to other research seeking to analyze and prevent interactions between proteins. This research is also unique because it primarily utilizes structural approaches and does not require extensive knowledge of any proteins. This approach can be particularly viable when newly discovered molecules are found to be harmful, as it provides a method to mitigate their effects by preventing specific interactions.

## Acknowledgement

This work is also partially supported by a start-up grant from Albert Einstein College of Medicine. Computational support was provided by Albert Einstein College of Medicine High Performance Computing Center.

## Author Contributions

V.L. and Y.W. designed research; V. L. performed research; V. L. analyzed data; V.L. and Y.W. wrote the paper.

## References

1. Ansarullah Jain, C., Far, F.F. et al. Inceptor counteracts insulin signaling in β-cells to control glycemia. Nature 590, 326–331 (2021).

2. McAulay, V., Deary, I.J. and Frier, B.M. (2001), Symptoms of hypoglycaemia in people with diabetes. Diabetic Medicine, 18:690–705. 10.1046/j.1464-5491.2001.00620.x

3. Lizcano, J. M., & Alessi, D. R. (2002). The insulin signalling pathway. Current Biology, 12(7), R236–R238. doi:10.1016/S0960-9822(02)00777-7

4. Bryant, P., Pozzati, G., Zhu, W. et al. Predicting the structure of large protein complexes using AlphaFold and Monte Carlo tree search. Nat Commun 13, 6028 (2022).

5. J;, Z. H. H. (2018a). Findsitecomb2.0: A new approach for virtual ligand screening of proteins and virtual target screening of biomolecules. Journal of chemical information and modeling. https://pubmed.ncbi.nlm.nih.gov/30278128/

6. Labbé CM, Rey J, Lagorce D, Vavruša M, Becot J, Sperandio O, Villoutreix BO, Tufféry P, Miteva MA. MTiOpenScreen: a web server for structure-based virtual screening. Nucleic Acids Res. 2015 Jul 1;43(W1): W448–54. doi: 10.1093/nar/gkv306. Epub 2015 Apr 8. PMID: 25855812; PMCID: PMC4489289.

7. Xiaoqiang Huang, Robin Pearce, Yang Zhang. FASPR: an open-source tool for fast and accurate protein side-chain packing. Bioinformatics (2020) 36: 3758–3765.

8. Xue L., Rodrigues J., Kastritis P., Bonvin A.M.J.J.*, Vangone A.*, “PRODIGY: a web-server for predicting the binding affinity in protein-protein complexes”, Bioinformatics, doi:10.1093/bioinformatics/btw514 (2016).

9. Feyza Maden, S., Sezer, S., & Ece Acuner, S. (2023). Fundamentals of Molecular Docking and Comparative Analysis of Protein–Small-Molecule Docking Approaches. IntechOpen. doi: 10.5772/intechopen.105815

10. U.S. National Library of Medicine. (2014, December 1). Type A insulin resistance syndrome: Medlineplus Genetics. MedlinePlus. https://medlineplus.gov/genetics/condition/type-a-insulin-resistance-syndrome/#:~:text=They%20develop%20cysts%20on%20the,thick%2C%20dark%2C%20and%20velvety.

11. Patterson, J. H., & Watkins, L. (2006, February 18). Leprechaunism in a male infant. The Journal of Pediatrics. https://www.sciencedirect.com/science/article/abs/pii/S0022347662801000

12. Bathi, R.J., Parveen, S., Mutalik, S. et al. Rabson-Mendenhall syndrome: two case reports and a brief review of the literature. Odontology 98, 89–96 (2010). 10.1007/s10266-009-0106-7

13. Li, M., Chi, X., Wang, Y. et al. Trends in insulin resistance: insights into mechanisms and therapeutic strategy. Sig Transduct Target Ther 7, 216 (2022).

14. Knox C, Wilson M, Klinger CM, et al. DrugBank 6.0: the DrugBank Knowledgebase for 2024. Nucleic Acids Res. 2024 Jan 5;52(D1): D1265–D1275. doi: 10.1093/nar/gkad976.

